# The relationships between genetic ancestry, somatic mutation frequency, and histologic subtypes in high-grade endometrial cancer

**DOI:** 10.1101/2023.07.26.550722

**Authors:** Ryan Bremseth-Vining, Victor Borda, Douglas Craig, Julie J. Ruterbusch, Julie Boerner, Juliana Fucinari, Rouba Ali-Fehmi, Mohamed Elshaikh, Hassan Abdallah, G. Larry Maxwell, Kathleen M. Darcy, Gregory Dyson, Thomas Conrads, Nicholas W. Bateman, Michele L. Cote, Timothy D. O’Connor

**Affiliations:** Institute for Genome Sciences, University of Maryland School of Medicine, Baltimore, MD 21201, USA; Wayne State University School of Medicine, Department of Oncology, Detroit, MI 48201, USA; Wayne State University School of Medicine, Department of Pathology, Detroit, MI, USA; Karmanos Cancer Institute, Tumor Biology and Microenvironment Program, Detroit, MI, USA; Henry Ford Hospital, Department of Radiation Oncology, Detroit, MI, USA; Women’s Health Integrated Research Center, Annandale, VA; Women’s Health Integrated Research Center, Falls Church, VA; Simon Comprehensive Cancer Center, Indiana University; Program in Health Equity and Population Health, University of Maryland School of Medicine, Baltimore, MD 21201, USA; Program in Personalized Genomic Medicine, University of Maryland School of Medicine, Baltimore, MD 21201, USA; Department of Medicine, University of Maryland School of Medicine, Baltimore, MD 21201, USA

**Keywords:** Local ancestry, somatic mutations, diplotype, endometrioid carcinoma, serous carcinoma, endometrial cancer, ethnicity, cancer epidemiology.

## Abstract

High-grade endometrial cancer, like numerous other cancer types, exhibits clear racial disparities in the United States for both the incidence and outcomes of the disease. While institutional factors are likely the primary contributor to these disparities, other underlying causes cannot be ignored (i.e., molecular, genetic, and histopathologic factors). This study seeks to interrogate the role that germline genetic influences, specifically genetic ancestry, may play in contributing to characteristics of high-grade endometrial cancer. This is mainly accomplished by examining the relationship between local ancestry inferences and somatic mutation frequency as well as histologic subtypes. An association between clinical characteristics and patient survival was also interrogated, and while global ancestry was seen to have no significant effect, tumor mutation burden (TMB) did impact patient survival. Here, we identify associations between local ancestry segments on chromosomes 1 and 14 and an increased TMB in self-described (SD) Black patients. We also highlight a complex relationship between heterozygous ancestry combinations within genomic regions (i.e., [European/African] vs. [African/African]) and an increase in local somatic mutation frequency.

Furthermore, we explore the relationship between local ancestry and histologic subtype. We identify one region (chr9q32) wherein the African/European local ancestry diplotype was associated with a higher incidence of serous carcinoma. We also underline a difference in somatic mutation frequency between endometrioid and serous carcinoma. While highly exploratory, these findings begin to characterize the complex relationship between genetic ancestry and characteristics of high-grade endometrial cancer, which may impact patient survival.

## Introduction

Due to advances in medical technology, targeted therapies, and behavioral changes, many cancer types have shown decreased incidence over the last 20 years. Endometrial cancer incidence, however, has risen by about 1% per year over that same period with recent stability (1), even after adjustment for hysterectomy prevalence (2). This rise in overall endometrial cancer risk is paired with exhibited disparities in the incidence and outcomes of the disease between racial and ethnic groups (3,4). Specifically, for self-described (SD) Black women in the United States, a higher incidence of endometrial cancer is accompanied by an increased presentation of highly aggressive histological subtypes (5,6). Socioeconomic factors and comorbidities that disproportionately affect Black individuals play a significant role in creating disparities in the incidence of endometrial cancer but do not explain them entirely (7,8).

Consequently, the mortality rate in SD-Black women is more than twice the rate observed in SD-white women after an endometrial cancer diagnosis. Even in instances when these individuals have equal access to healthcare, the differences in outcomes are still observed. This suggests that the observed racial disparities in endometrial cancer outcomes are likely related to numerous other extrinsic and intrinsic factors (9).

Molecular, histopathologic, and genetic elements contribute to these disparities in patient outcomes and treatment response. Tumor mutational burden and gene-specific mutations have been demonstrated as effective indicators of treatment response and prognostic outcomes in endometrial cancer (10–12). Moreover, germline genetic ancestry has been shown to associate with targetable mutations and prognostic biomarkers, including tumor mutation burden, in various other cancer types (13,14). Specifically for endometrial cancer, increased levels of genome-wide African ancestry have been suggested to associate with decreased patient survival after stratifying by stage and race (15). Here, we extend this observation by examining the effect of genome-wide ancestry and relevant clinical characteristics on the survival of patients in our study population of women with high-grade endometrial cancer. Although these genome-wide ancestry estimates can inform us about population stratification and its effect on survival, a more subtle analysis of ancestry (i.e., changes of continental ancestry across the genome) is needed to understand the impact of genetic ancestry on differences in the characteristics/outcomes of endometrial cancer and how they relate to patient survival.

To that end, we explored how the ancestral background, in terms of the ancestry of genomic regions, correlates with the mutational landscape and histopathological characteristics in women with high-grade endometrial cancer. To address this question, we inferred genome-wide and local ancestry backgrounds from germline whole-exome data of 285 women (including 164 SD-Black individuals) obtained from two academic medical centers in Detriot, Michigan. By coupling ancestry inferences with somatic mutation burden and clinical data (16), we performed regression modeling to determine its relationship with the somatic mutation landscape and histopathological subtype. We also interrogated the effect of the combination of local ancestry haplotypes of differing ancestries on somatic mutation burden and histologic type.

## Methods

### Study samples

Potential cases were identified from observational studies at Henry Ford Health System (HFHS) and Karmanos Cancer Institute (KCI). Data from the Metropolitan Detroit Cancer Surveillance System (MDCSS) was also used. Eligible cases included non-Hispanic black and white women, ages 21–79 at diagnosis, who underwent a hysterectomy at HFHS or KCI at the time of diagnosis between 1997-2016. Only those with the International Federation of Gynecology and Obstetrics (FIGO) 2009 grade III at the time of diagnosis were included. Additionally, as access to tissue was required, only women with FIGO stages I-III tumors were included. Detailed clinical and pathological data, including patient age of diagnosis, body mass index (BMI), histological type, and progression-free survival in months, were collected.

### Endometrial tissue processing

Representative hematoxylin and eosin (H&E) stained thin sections from each formalin-fixed, paraffin-embedded endometrial cancer specimen were pathologically evaluated to assess tumor cellularity and necrosis. Any samples exhibiting >20% necrosis or <25% tumor cellularity were excluded. All tissue specimens were thin-sectioned (8 mm) onto polyethylene naphthalene (PEN) membrane slides and lightly stained with H&E. Specimens that were determined to be >70% tumor cellularity were manually scraped to harvest tissue. Specimens that were <70% tumor cellularity were selectively harvested by laser microdissection to enrich the final sample to >70% in tumor epithelium. Normal adjacent tissue was collected by manual scraping to generate a germline control DNA sample.

### DNA extraction and Exome sequencing

DNA Isolation was performed using the QiAamp DNA Micro Kit (Qiagen Sciences LLC, Germantown, MD) according to the manufacturer’s protocol. DNA was eluted after 10 min incubations with 40 μL and 100 μL of Buffer AE and 100 μL of nuclease-free water, respectively, and reduced to 40 μL by vacuum centrifugation. Initial quantity and 260/280 purity readings were assessed spectrophotometrically (Nanodrop 2000 Spectrophotometer, Thermo Fisher Scientific). Final quantities were determined using Qubit DNA HS and BR kits (Thermo Fisher). A total amount of 1.0 μg genomic DNA per sample was used as input material for the DNA library preparation. Sequencing libraries were generated using the Agilent SureSelect Human All Exon kit (Agilent Technologies, CA, USA) following the manufacturer’s recommendations, and index codes were added to each sample. Briefly, DNA was fragmented by a hydrodynamic shearing system (Covaris, Massachusetts, USA) to generate 180-280 bp fragments, overhangs were converted into blunt ends via exonuclease/polymerase activities, and enzymes were removed. After the adenylation of DNA fragment 3’ ends, adapter oligonucleotides were ligated. PCR selectively enriched DNA fragments with ligated adapter molecules on both ends. After PCR, libraries were hybridized with a biotin-labeled probe, after which streptomycin-coated magnetic beads were used to capture exons. PCR-enriched captured libraries to add index tags in preparation for hybridization. Products were purified using AMPure XP system (Beckman Coulter, Beverly, USA) and quantified using the Agilent high-sensitivity DNA assay on the Agilent Bioanalyzer 2100 system. Next-generation sequencing was conducted with the NovaSeq 6000 platform (Illumina).

### Quality control of sequencing data

We employed multiple approaches to assess the quality of the data and to identify if any of the patient pairs needed to be excluded from the downstream analyses. Of the resulting 19 patients excluded, the reasoning involved either a mixed diagnosis, a stage 4 case, or a failed whole exome sequencing QC check. After applying these filters, 285 [CML2] usable genomic sample pairs that we are confident are from the same patient and of sufficient quality remain for subsequent analyses.

For somatic variant calling, filtering the Mutect2 identified variants was necessary due to the overabundance of C>T (and A>G) SNPs observed, which are a feature of FFPE contamination. Aggressive filtering steps were utilized, and 63% of the initially passed variants (842,456 out of the 1,351,199 variants over the 285 samples) were eventually excluded. We define functional variants using the results from ANNOVAR as those variants that were splicing, frameshift insertion, frameshift deletion, frameshift substitution, nonframeshift insertion, nonframeshift deletion, nonframeshift substitution, stopgain, stoploss; or had at least 50% of the functional callers (including SIFT, Polyphen HDIV, Polyphen HVAR, LRT, Mutation Taster, Mutation Assessor, RadialSVM) identified the site as deleterious.

For our germline whole-exome data, we removed variants with missing data above 5% and out of Hardy Weinberg equilibrium using plink2 (17) (--geno 0.05 and --hwe 10e-6, respectively) and kept 285 individuals with 611,625 variants. Our data included 164 SD-Black and 121 SD-White. To control for cryptic relatedness, we ran REAP (18) to determine the kinship coefficient among pairs of individuals. After relatedness analysis, we found two pairs of individuals with a first-degree relationship. We removed one of the individuals of each pair corresponding to SD-White individuals.

### Genome-wide ancestries, Mutation Burden, and Survival

Population structure analyses were performed through principal component analysis (PCA) and genome-wide ancestry proportions estimated with RFMIX (19) on germline data. We ran PCA using SNPRelate (20) for our entire dataset (SD-Black and SD-White individuals) after filtering by MAF and removing highly linked SNPs (r^2^>0.4).

Kaplan Meier curves were plotted, and Aalen’s Additive regression was fit to the data (to account for non-static effects of covariates over time) using the R package Survival. Genome-wide ancestry proportions were included in this regression, as were relevant demographic and clinical variables, including tumor mutation burden. These regressions were performed for the entire study group, as well as for groups stratified by tumor histology and stage.

We perform a negative binomial regression to model the relationship between each of the four genome-wide ancestry proportions and total somatic mutation burden (TMB). We adjusted the regression considering the age of diagnosis, body mass index (BMI), and histological subtype of tumor (i.e., clear, mixed, endometrioid, serous carcinoma).

### Phasing and Local Ancestry Inferences

We inferred the haplotype phase for our data using SHAPEIT 4 (21) with TOPMed (22) data as a reference panel. This panel includes the high-coverage version of the 1000 Genomes Project (23). After phasing, we performed a local ancestry inference (LAI) using RFMix ver2 (19) with two expectation-Maximization runs, eight generations since admixture, and a subset (n=1,465 individuals) of the TOPMed panel encompassing four ancestry groups: African, European, East Asian, and Indigenous Americans. The African ancestry was represented by 405 individuals from LWK, YRI, GWD, and MSL populations. European ancestry was represented by 404 individuals from GBR, TSI, IBS, and CEU. East Asian ancestry was represented by 312 individuals from CHB, CHS, and JPT. The Indigenous American ancestry included 344 Latin American individuals with a high proportion of this ancestry (>80%) from the 1000 Genomes Project (PEL and MXL) and the TOPMED Project. Considering that our Indigenous American references are admixed individuals, we included the rfmix flag--reanalyze-reference. For our association analyses, we identified a genome-wide significance threshold for our admixture mapping analyses using STEAM (24) and considering eight generations since admixture and default parameters.

### Relationship between Somatic Mutations Abundance and Ancestry Haplotype Count

After LAI, we count the number of ancestry haplotypes (i.e., the presence of 0, 1, or 2 haplotypes for a specific ancestry in a genomic region) for each individual for each genomic window identified by the conditional random fields approach of RFMIX. Considering the genetic coordinates of the ancestry windows, we determine the abundance of somatic mutations in each window for each individual. We also compute the frequency of nonsynonymous mutations across the genome (i.e., TMB, or nonsynonymous mutations/Mb). To examine the relationship between Ancestry counts and the number of somatic mutations in each local ancestry block as well as the burden of nonsynonymous mutations across the genome, we ran a negative binomial regression using the R package MASS. This regression was adjusted by the “Age at diagnosis” as well as the first four principal components (PCs). A linear approximation was substituted if the negative binomial model failed to converge at a value of theta for a local ancestry region.

### Distribution of Density of Somatic Variants within ancestry-diplotypes

Considering that SD-Black individuals in our sample are admixed compared to SD-white, we focused our subsequent analyses only on the SD-Black (n=164). Our somatic mutation data for each individual contains a significant proportion of variants that are not present in common with our germline data. For this reason, we were not able to determine the haplotype phase or local ancestry haplotype for somatic variants. To overcome this problem and determine the relationship between the density of somatic mutations and Local ancestry, we introduce the concept of diplotype at the individual level. We call the diplotype of a genetic region the combination of local ancestry haplotypes for a region determined by RFMIX. Considering that SD-Black have predominantly European and African ancestries, we focused on three diplotype combinations: Eur/Eur, Eur/Afr, and Afr/Afr.

To compare if a specific diplotype harbors more somatic mutations than the other in each individual, we determined the genome-wide average of somatic mutations per diplotype and divided it by the total number of base pairs of the diplotype per individual. We multiplied by 1,000,000 to have an index by Mb. After getting the per-individual somatic/diplotype index (piSD index), we compared two density values of a specific diplotype of an individual only if the individual harbors a certain amount of the diplotype (10, 20, and 30Mb). After filtering, we plot the difference between the piSD index of different haplotypes in an individual. We estimate if the difference between indexes was different from zero using a Wilcoxon test. We stratified our data based on the histopathological phenotype (i.e., endometrioid and serous adenocarcinoma). We performed the comparison of densities in each histopathological type.

### Ancestry Diplotypes, Density of Somatic variants, and Heterozygosity

To explore other possible drivers of somatic variants, we determined (i) the relationship between ancestry diplotypes and heterozygosity and (ii) the relationship between the density of somatic variants with heterozygosity. For each local ancestry window, we estimated the density of somatic mutations and the inbreeding coefficient (F statistic) per individual in the germline data. The inbreeding coefficient was estimated using vcftools that use the equation F = (O − E) / (N − E), where O corresponds to the observed number of homozygotes, E is the expected number of homozygotes (given population allele frequency), and N is the total number of genotyped loci in the specific window. Negative F values indicate an excess of heterozygosity for the corresponding window. To examine the relationship between ancestry diplotypes and F values, we compare the distribution of F values between pairs of diplotypes using a Wilcoxon test.

### Relationship Between African Ancestry and Histological Characteristics

A Firth regression was used to model the association between African and European ancestry haplotype counts and histological subtypes (endometrioid and serous adenocarcinoma). This regression was chosen to accommodate our small sample size for this subgroup in which the SD-Black individuals with mixed histological subtypes were excluded (n = 135). Also, we performed a Firth regression on the SNP variants in each of the genomic regions that resulted significantly in the latter regression modeling. Briefly, we model the histopathological outcome considering genotype information and adjusting by age and the first five PCs. Furthermore, we looked for differences in the density of rare variants and the histopathological phenotype using SKAT (25).

### Code availability

Scripts to perform all statistical analyses and plotting results are available at: https://github.com/umb-oconnorgroup/Endometrial-Cancer

### Data availability statement

The data generated in this study are available upon request from the corresponding author.

## Results

### Global Ancestry, somatic mutations, and survival outcome

Our dataset included SD-Black (n=164) and SD-White individuals (n=121) with extensive clinical data, including patient age of diagnosis, body mass index (BMI), histological type, and progression-free survival in months. To explore the genetic composition and population structure, we performed genetic clustering (Figure 1A) and PCA (Supplementary Fig 1) on germline whole-exome data. Both analyses showed a tight relationship between self-described race and genetic ancestry. The SD-Black individuals showed a more heterogeneous ancestry composition represented by a significant proportion of European and African ancestries in contrast to the SD-white individuals.

**Figure 1.**
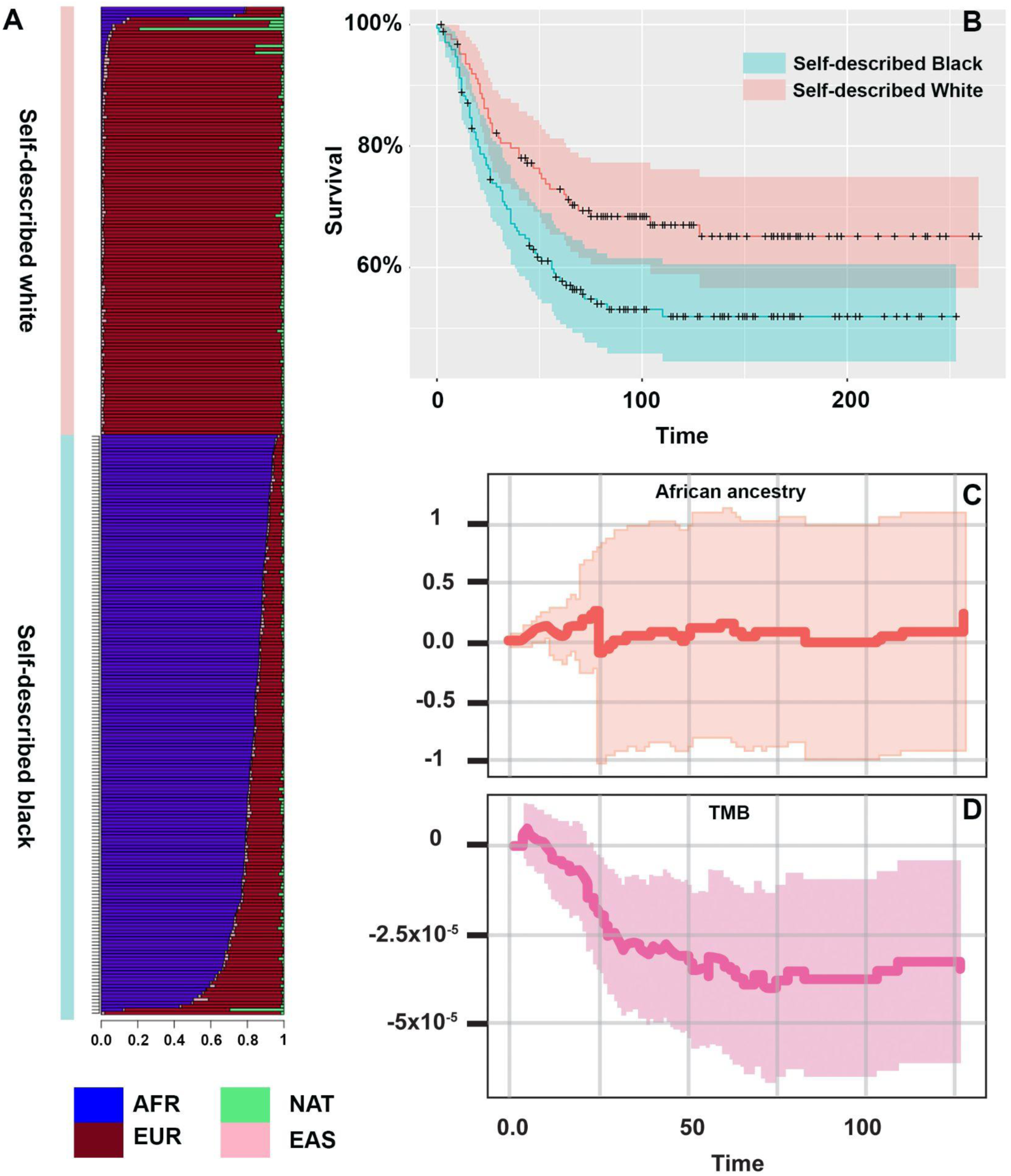
Ancestry composition and survival analysis of SD-Black and SD-White samples with endometrial cancer. (A) Global ancestry proportions inferred with RFMIX ver 2. The plot includes SD-white (brown bar) and SD-Black (purple bar). (B) Kaplan Meier curves show survival over time (months) and stratified by self-described race. SD-Black individuals are represented by the curve highlighted in blue, and the SD-white individuals are represented by the curve highlighted in red. (C) and (D) Plots show the effect of AFR ancestry proportions and Total mutation burden (TMB) on the hazard ratio over time, respectively, as computed by Aalen’s Additive regression, including the entire study group. Reference populations for ancestry: AFR: African ancestry, EUR: European ancestry, NAT: Indigenous American ancestry, EAS: East Asian ancestry.

We addressed the relationship between survival outcome (including progression-free survival) with (1) global African ancestry and (2) tumor mutation burden (TMB). First, to follow up on the results that Rocconi et al. (2016) described, wherein the authors identified a trend between increased African ancestry proportions and decreased survival, we examined the association via survival analysis. In contrast to what was described by Rocconi and colleagues, we observed no association between African ancestry proportions and differential survival for our entire dataset nor when restricting the analysis to the SD-Black individuals (Figure 1B and C).

Second, we test for a relationship between survival and TMB, or the burden of nonsynonymous mutations, which is an important predictor for cancer treatment since tumors with high TMB have a better response to specific treatments (26,27). We obtained somatic mutation data from endometrial cancer tissue corresponding to the same individuals for germline data. We observed a significant association between greater survival and a higher TMB (p-value = 7.32 x 10^-3^, Figure 1D). This association also was observed when considering progression-free survival in these same patients. TMB was still significantly associated with better progression-free survival (p = 2.23 x 10^-3^) when considering all of the patients within this study group, and African and European ancestry proportions still showed little to no association.

Third, we explored the relationship between TMB and the genome-wide ancestry proportions through regression modeling adjusting by the age of diagnosis, body mass index, and histological subtype for the entire dataset and self-described black individuals. We found no association (p-value> 0.1) between TMB with any of the four ancestries (African, European, Indigenous American, and East Asian). Considering that SD-White individuals have predominantly European ancestry (mean _European ancestry_= 93%), contrasting with SD-Black (mean _European ancestry_= 16%), we restricted the subsequent analyses to SD-Black individuals.

### Chromosomal Ancestry background and burden of somatic mutations

Our latter results showed that genome-wide ancestry does not explain the observed survival disparities nor the density of somatic mutations within these patients. However, other levels of ancestry still need to be explored. We investigated the relationship between the density of somatic mutation and ancestry changes along individual chromosomes (i.e., local ancestry) through two approaches: admixture mapping and ancestry diplotype analyses. We infer the local ancestry patterns in our germline whole-exome data using RFMIX ver. 2 (19). Using STEAM (24), we identified a genome-wide significance threshold of 1.046 x 10^-5^ for our admixture mapping analyses.

We performed admixture mapping analyses to identify (i) the relationship between local ancestry counts and the local frequency of somatic mutations (i.e., within the same genomic region) as well as (ii) the relationship between the same local ancestry counts with the genome-wide frequency of all somatic mutations, and tumor mutation burden (i.e., the frequency of nonsynonymous mutations across the genome). Considering the lower genome-wide proportion of Indigenous American and East Asian ancestries, we focused subsequent analyses only on African and European ancestries. We included the first four PCs and patient age at diagnosis for each model as covariates. We found no region reaching the genome-wide significance threshold for the models regarding African ancestry haplotypes and local mutation frequency, as well as total somatic mutations across the genome (Supplementary Fig. 2). For the model regarding African local ancestry segments and TMB; we observed three regions; two contiguous regions on chromosome 1 and one region on chromosome 14 reaching genome-wide significance (p-value < 1.046 x 10^-5^; Figure 2, Supplementary Fig 2B, and Supplementary Table 2).

**Figure 2.**
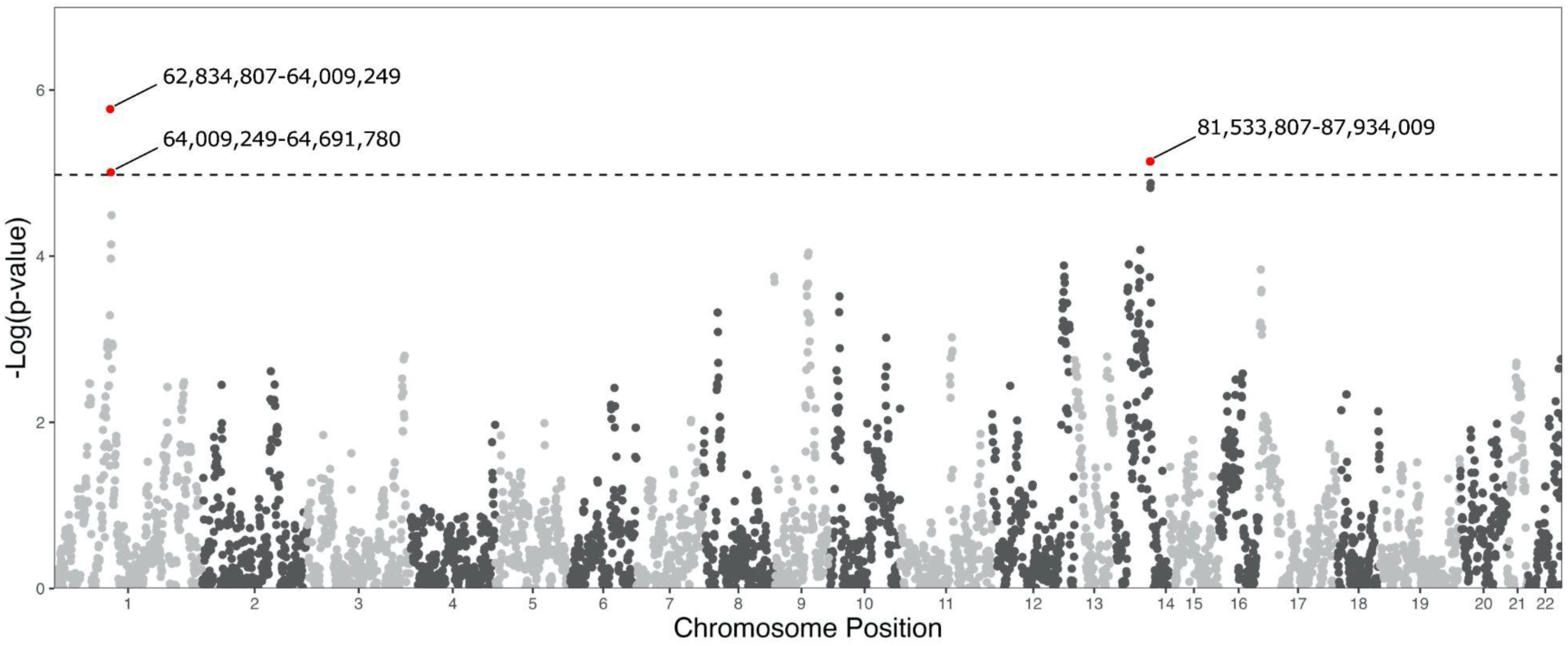
Effects of local ancestry enrichment on the burden of nonsynonymous mutations. Manhattan plot of the genome-wide distribution of p-values resulting from a model investigating the relationship between the count of African local ancestry haplotypes per local ancestry region and the total burden of nonsynonymous mutations across the genome. We used age at diagnosis and principal components 1-4 as covariates in this model. Genomic regions that exceeded our genome-wide significance threshold are highlighted, and the position of those regions is listed.

These regions on chromosome 1 contain loci that have been associated with the incidence of ovarian cancer as well as breast cancer, and the chromosome 14 region highlighted in this analysis includes a gene associated with meningioma progression (28–30). These associations, however, have seemingly not been previously interrogated in the context of individuals with appreciable levels of African ancestry. These former analyses were also conducted for European haplotypes, but none of these models identified any statistically significant correlations (Supplementary Fig. 3).

One major issue in determining the ancestry background for somatic mutations in a specific genomic region is the inability to determine the phase of somatic variants since most of these variants are not represented in any reference panel. To overcome this problem, we organized our local ancestry data into ancestry diplotypes. We define an ancestry diplotype as the combination of the local ancestry inferences of both haplotypes in each genomic region for each individual (e.g., homozygous African [Afr/Afr], heterozygous African and European [Eur/Afr], etc.) (Figure 3 A). After identifying the ancestry diplotype, we estimated the density of somatic mutations within the same genomic region. Finally, we computed a “*Per-individual somatic/diplotype index*” (piSD index) as the average of observed somatic mutations per Mb of an ancestry diplotype per individual.

**Figure 3.**
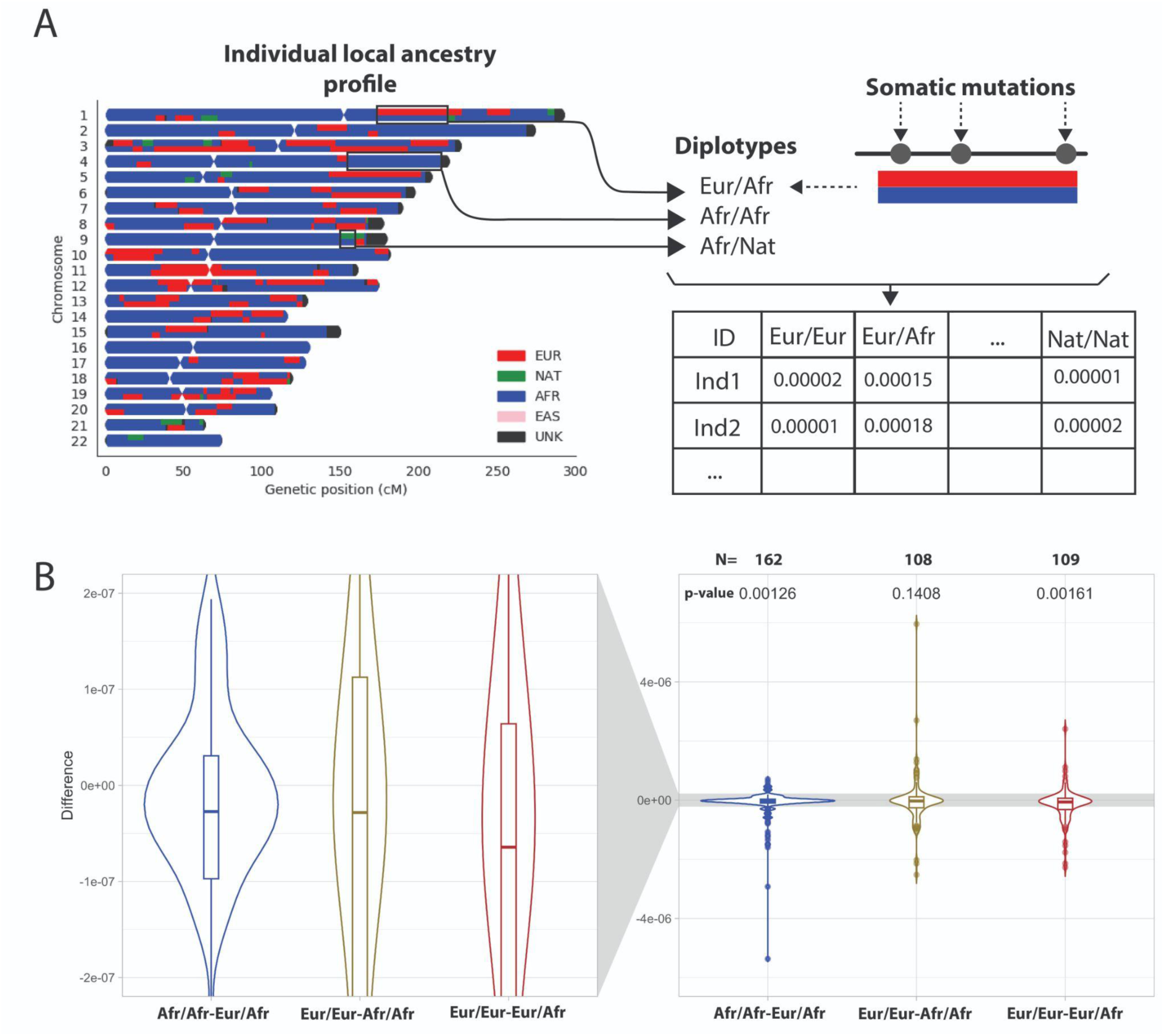
Relationship between Ancestral background of genomic regions and density of somatic mutations. A) We calculated the Per-individual somatic/diplotype index (piSD index) by identifying the number of somatic mutations per Mb per diplotype (i.e., Eur/Afr, Afr/Afr, Afr/Nat) B) Frequency distribution of somatic mutations per Local ancestry interval per diplotype. B) Violin plots showing the distribution of the differences between pairs of piSD indexes. Each point represents the difference between diplotype rates in the same individual. We restricted the estimation of the differences to individuals harboring at least 30Mb of each diplotype being compared. Supplementary Fig 4 shows the distribution of differences for individuals harboring at least 20 and 10 Mb. Sample size and p-values are shown at the top of the plots.

To determine whether an ancestry diplotype is harboring more somatic mutations than another diplotype, we estimated the difference between the piSD index of two ancestry diplotypes for each individual (e.g., piSD index _Eur/Eur_ - piSD index _Eur/Afr_) and determined if the median of the distribution of differences was significantly different from zero. Our pairwise comparisons showed that the differences between the pairs Eur/Eur - Eur/Afr and Afr/Afr-Eur/Afr are significantly negative values (Figure 3B and Supplementary Fig 4; p-value_Eur/Eur-Eur/Afr_=0.00161; p-value_Afr/Afr-Eur/Afr_=0.00126). This indicates heterozygous Eur/Afr regions harbored a higher density of somatic mutations than homozygous European and African regions.

It is suggested that homologous chromosomal regions that differ in sequence, harboring heterozygous sites, have a high rate of *de novo* mutations, potentially due to incorrect recruitment of mismatch repair mechanisms (31). We tested whether a similar trend could explain the relationship between Eur/Afr and somatic mutations. Under this hypothesis and considering the heterozygous Eur/Afr diplotype harboring the higher genome-wide rate of somatic mutations, we tested if: i) Eur/Afr diplotypes also have higher levels of heterozygosity compared to the homozygous diplotypes (i.e., Afr/Afr and Eur/Eur) and ii) if a higher level of heterozygosity is associated with a higher density of somatic mutations. We estimated heterozygosity by calculating the inbreeding coefficient (F), whose negative values indicate an excess of heterozygosity. We estimated F for each of the intervals of local ancestry per individual. We found that heterozygous Eur/Afr regions (F mean = 0.02745) have a lower F value compared to the homozygous Eur/Eur (F mean = 0.31191) and Afr/Afr (F mean = 0.03726) regions (Supplementary Fig 5). However, we did not find any significant relationship between the density of somatic mutations and the inbreeding coefficient (Supplementary Fig 6). This suggests that heterozygosity alone may not be sufficient to account for the differences in piSD index observed between diplotypes.

### Chromosomal Ancestry backgrounds and histopathological characteristics

Endometrial cancer is often classified based on histopathological characteristics. These histological types include endometrioid, clear cell, and serous carcinoma, the latter being the most aggressive type (32). The clinical data available for our dataset allowed us to explore the relationship between the ancestral background and somatic mutation frequency by histological type. Two histological types, endometrioid (n_SD-Black_=51) and serous (n_SD-Black_=85) carcinoma, were the most frequent in our data. By stratifying our data in these two types and exploring the piSD index, we found different patterns within the SD-Black patients for these two histology types (Figure 4A and 4B and Supplementary Fig 7). For the endometrioid histologic type, heterozygous Eur/Afr had a significantly higher rate of somatic mutations when compared to homozygous Afr, and the serous type shows a significant value when Eur/Afr is compared to homozygous Eur. Interestingly, by examining the somatic rate across histological types and functional types of variants (i.e., nonsynonymous, INDELS, transitions), the endometrioid type showed a higher rate of somatic mutations compared to the serous type independent of the ancestry background (Figure 4C and 4D and Supplementary Fig 8 and 9).

**Figure 4.**
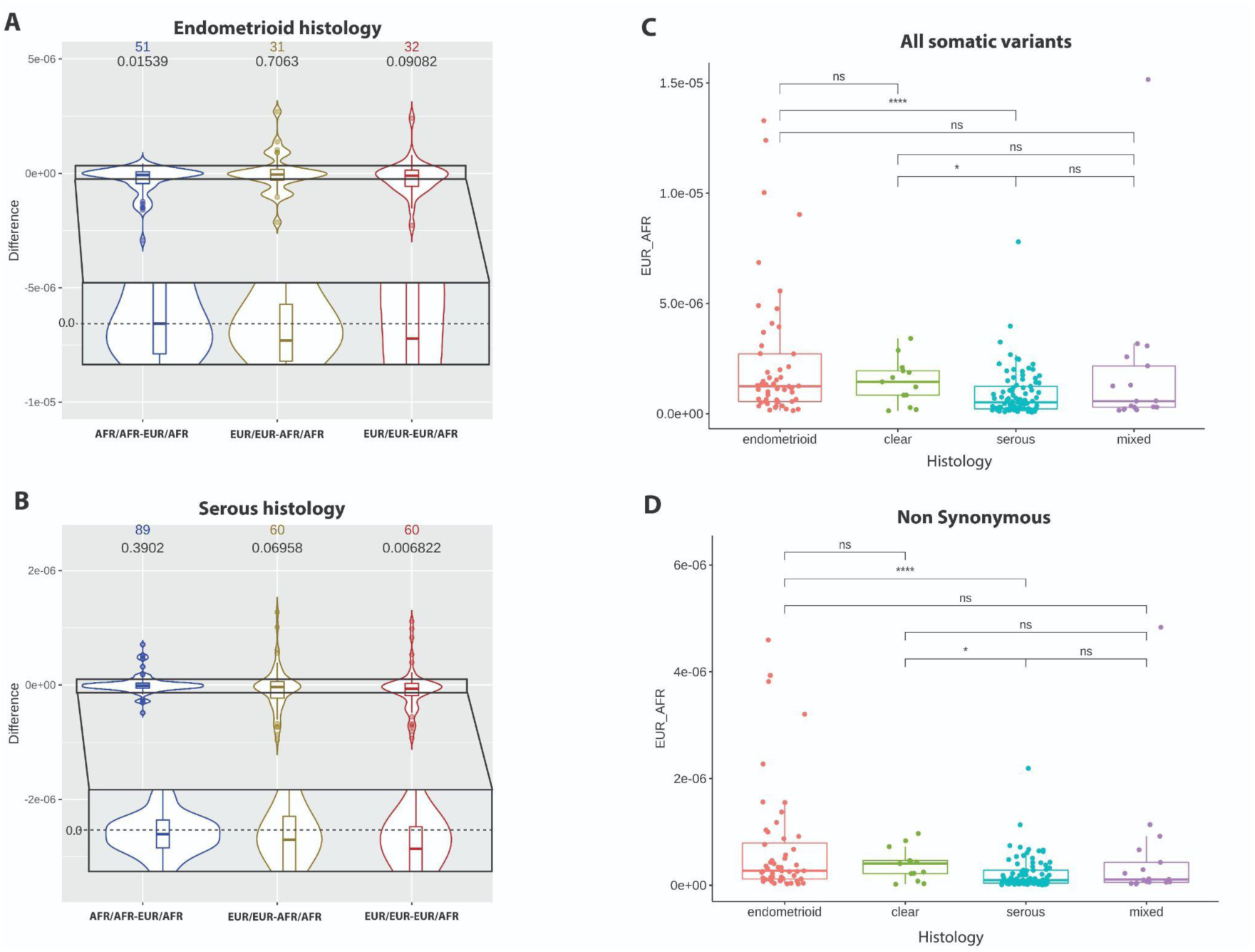
Relationship between Ancestral background of genomic regions and density of somatic mutations per Histological type. A) Violin plots showing the distribution of the differences between pairs of piSD indexes in individuals with endometrioid histology subtype. B) Violin plots showing the distribution of the differences between pairs of piSD indexes in individuals with serous histology subtype. C) Average mutation burden per Mb observed in EUR/AFR diplotype per individual stratified per histological type (i.e., endometrioid, clear, serous, mixed). D) Average nonsynonymous mutation burden per Mb observed in EUR/AFR diplotype per individual stratified per histological type (i.e., endometrioid, clear, serous, mixed). The significant difference between distributions is indicated on the top of the box plots. ns: non-significant, *: (0.01, 0.05], **: (0.001, 0.01], ***: [0, 0.001].

Next, we tested the association between histopathological type (i.e., endometrioid and serous) and local ancestry inferences (African and European haplotypes) across the genome in SD-Black individuals through admixture mapping. For this purpose, we applied a Firth regression model adjusting by age and the first five principal components. Using 1.046 x 10^-5^ as the significance threshold (24), we found one significant association for African ancestry (Figure 5A): 9q32 (OR: 0.112, 95% CI: 0.032-0.317, p-value=6.64×10^-6^). In contrast, this region did not reach the significance threshold for European ancestry (Figure 5B). By inspecting the frequency of histopathological type by the number of African counts for the 9q32 region (Table 1), we observed a contrasting proportion of individuals with endometrioid and serous carcinoma (∼1 endometrioid / 4 serous) when they harbor one copy of African or European haplotypes (Table 1). By verifying the local ancestry information for individuals with 1 African or 1 European copy, most of these segments harbor the heterozygous Eur/Afr diplotype. Our results suggest that having an Afr/Afr diplotype is associated with a similar occurrence of endometrioid and serous carcinoma, but having a heterozygous diplotype is associated with an overrepresentation of serous carcinoma.

**Figure 5.**
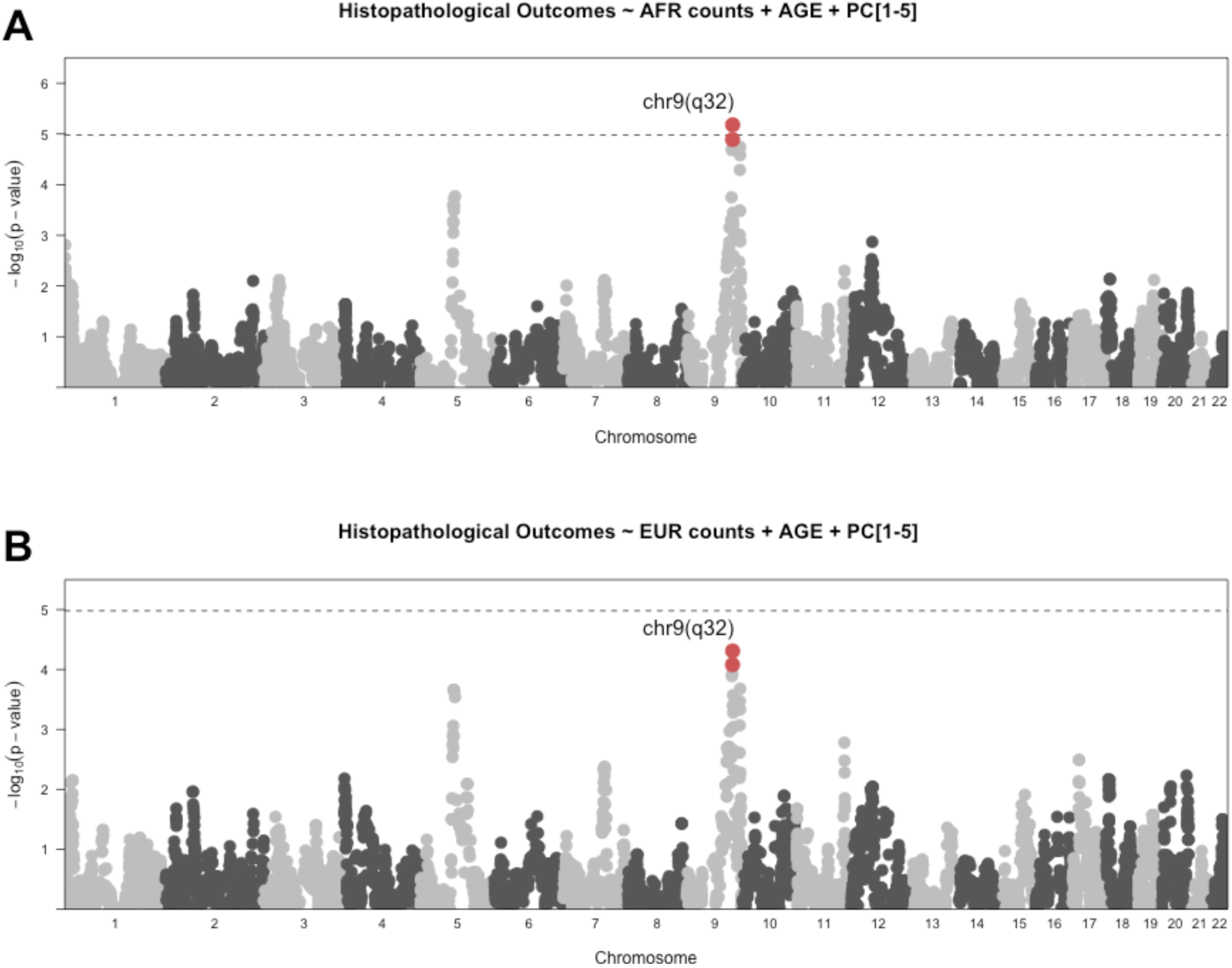
Manhattan plot showing admixture mapping peaks using Firth’s regressions with local ancestry inferences. One significant peak in 9q32 (chr9:112890610-113969108, Odd ratio: 0.112, 95% CI: 0.032-0.316, p-value=6.64×10^-6^). A) Admixture mapping for A) African and B) European counts.

**Table 1.**
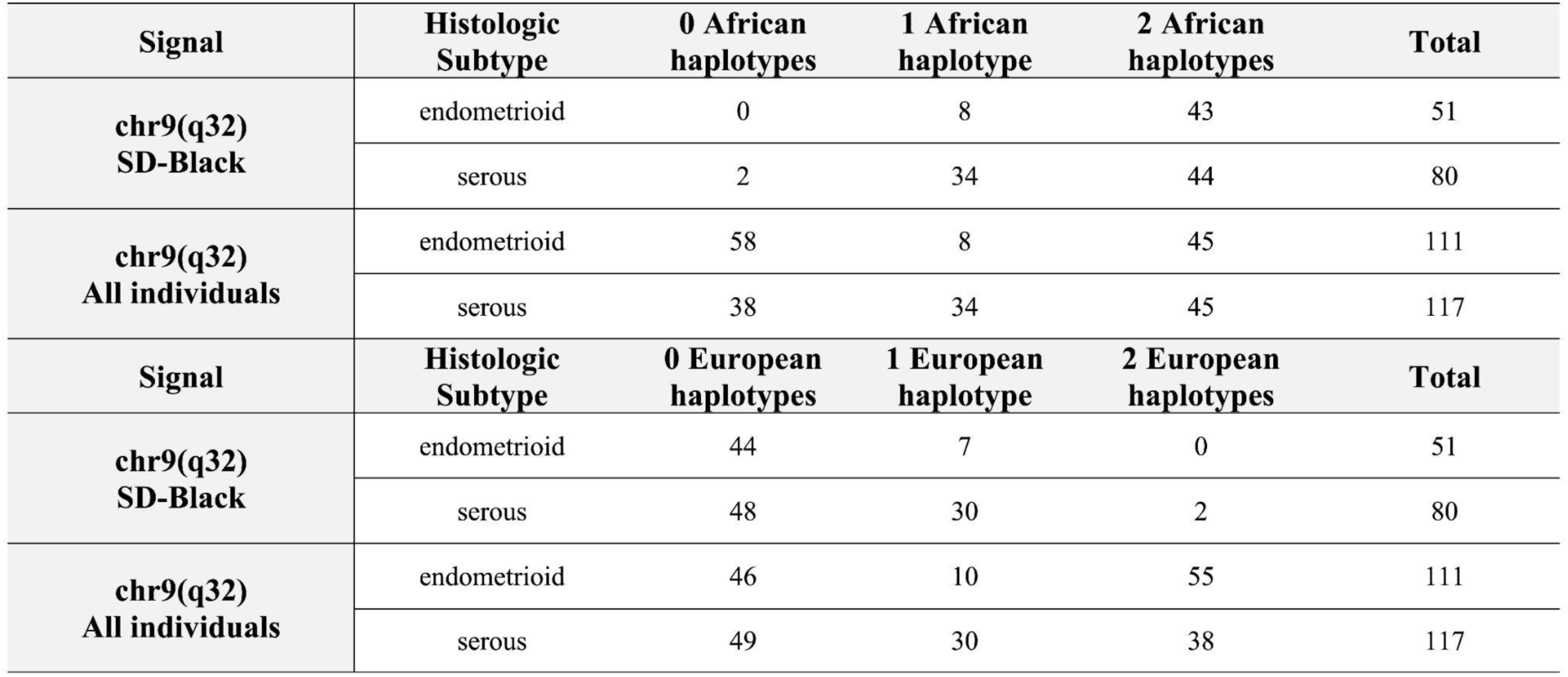
Contingency table showing the relationship between Ancestry haplotype counts in Local ancestry regions compared to the frequency of endometrioid and serous adenocarcinoma cases. Comparison of ancestry inference frequencies and histologic subtype incidence within the genomic region identified through Firth regression modeling (9q32: p = 6.64×10-6), shown for both SD-Black and all individuals.

| Signal | Histologic Subtype | 0 African haplotypes | 1 African haplotype | 2 African haplotypes | Total |
| --- | --- | --- | --- | --- | --- |
| chr9(q32)<br>SD-Black | endometrioid | 0 | 8 | 43 | 51 |
|  | serous | 2 | 34 | 44 | 80 |
| chr9(q32)<br>All individuals | endometrioid | 58 | 8 | 45 | 111 |
|  | serous | 38 | 34 | 45 | 117 |
| Signal | Histologic Subtype | 0 European haplotypes | 1 European haplotype | 2 European haplotypes | Total |
| chr9(q32)<br>SD-Black | endometrioid | 44 | 7 | 0 | 51 |
|  | serous | 48 | 30 | 2 | 80 |
| chr9(q32)<br>All individuals | endometrioid | 46 | 10 | 55 | 111 |
|  | serous | 49 | 30 | 38 | 117 |

To explore how this region may influence the incidence of the serous histologic subtype, we examined whether the heterozygous diplotype in chr9(q32) harbors highly differentiated variants or a higher density of missense and nonsense mutations compared to the homozygous. A regression analysis to identify associations between differentiated variants in this chromosome 9 region and histologic subtype did not return any significant results that could help further to explain our initial observation (Supplementary Fig 10). However, when we compare the accumulation of missense and nonsense variants in the same chromosome 9 region, we observed a significant difference in mutation frequency between the heterozygous and homozygous diplotype in the gene *POMT1* (p-value=0.0259).

## Discussion

We show how ancestry affects two endometrial cancer outcomes (i.e., the burden of somatic mutations and histological characteristics) by exploring patterns of global and local ancestry and integrating them with clinical data and somatic mutation data in 285 women. We have five main findings: (1) genome-wide African ancestry is not associated with differential survival; (2) three loci were identified wherein harboring African ancestry segments were associated with a higher tumor mutation burden; (3) genomic regions with haplotypes of contrasting ancestry backgrounds, which we define as heterozygous diplotypes, harbored a significantly higher genome-wide rate of somatic mutations compared to homozygous African and European genomic regions; (4) independent of the ancestry background of genomic regions, the endometrioid subtype showed a higher rate of somatic mutations compared to the serous subtype; (5) harboring a heterozygous diplotype at chromosome 9(q32) was associated with a higher incidence of serous carcinomas.

Racial differences have been observed in incidence and mortality rates between SD-Black and SD-White women with endometrial cancer (15,33). However, disparities are not exclusive to this type of cancer. For this reason, other risk factors like socioeconomic status and environmental factors have been postulated as drivers of these differences. Specifically for socioeconomic status, it was observed that it affects the evolution of endometrial cancer independent of racial disparities (34,35). Furthermore, to understand other factors that could trigger these disparities, genome-wide ancestry proportions were evaluated for association with outcomes in endometrial cancer. Previous associations have been suggested between increased levels of African ancestry across the genome and decreased survival (15). As our first main finding, we did not observe this trend in our study population, although we did observe a significant correlation between tumor mutation burden and survival within these patients.

A comparison of tumor evolution in different cancer types between African Americans (i.e., SD-Black) and European Americans (SD-White) has shown increased genomic instability in African Americans (36,37). These previous observations motivated us to explore subtle levels of admixture through local ancestry to identify the role of African ancestry in these disparities. Our second finding shows a positive correlation between the accumulation of African ancestry haplotypes in three regions across the genome and tumor mutation burden. This result, coupled with the association between greater survival and higher TMB, is important since a higher TMB has been associated with an improved patient response after specific immunotherapies (26,27). Moreover, the significant regions on chromosome 1 contain loci that have been associated with the incidence of ovarian cancer as well as breast cancer, and the chromosome 14 region highlighted in this analysis contains a gene associated with meningioma progression (28–30). These associations have not been previously interrogated in the context of African ancestry.

Our third finding is in the genome-wide context, as we observed that in genomic regions with a combination of European and African haplotypes (i.e., heterozygous diplotype), there was a greater accumulation of somatic mutations. This combination was common in SD-Black individuals but almost absent in SD-White individuals due to limited levels of admixture. We explored the hypothesis suggesting that heterozygosity may be correlated with aberrant recruitment of mismatch repair mechanisms leading to an increase in the local mutation rate (31). We found that an excess of heterozygosity (negative values of the inbreeding coefficient, F) was not correlated with a higher density of somatic mutations. On the other hand, the Eur/Afr diplotype showed the highest levels of heterozygosity within these patients. This suggests there are other mechanisms that could explain the different rate of somatic mutations in the Eur/Afr diplotype.

Our fourth and fifth findings explore the endometrial cancer histological types. SD-Black women in the United States have a higher incidence of aggressive subtypes (e.g., serous adenocarcinoma) compared to other racial/ethnic groups (38), which aligns with study population. We found that endometrioid tumors harbored more somatic mutations than serous tumors, and ancestry did not explain this difference. Interestingly, our admixture mapping results showed that for individuals harboring a heterozygous diplotype in chr9(32q) for our significant and suggestive results, the rate between endometrioid and serous becomes 1:4. This indicates that African ancestry could play a protective role at this locus.

Although our regression modeling using variants in these genomic regions resulted in non-significant signals, other subtle differences (e.g., accumulation of low-frequency variants) could generate these patterns.

A primary limitation is our small size (285 individuals, 164 SD-Black) and the low levels of admixture in the SD-White individuals, which does not allow us to extend our findings to this group. Nonetheless, our significant and consistent results offer new research opportunities for underrepresented groups with high-grade endometrial cancer. This study provides insights into how genetic ancestry associates with somatic mutations in high-grade endometrial cancer, but more work is needed to address these questions and disparities.

Future work requires a larger sample size to confirm observations. Currently, little can be concluded about the specific effects of African ancestry-derived alleles within these regions. However, the connection between African ancestry-associated alleles, TMB, and patient survival warrants further exploration. Local ancestry and admixture mapping can potentially identify consequential ancestry-associated alleles, even at small sample sizes, guiding future research.

Further efforts can examine associations between genetic ancestry and mutations in targetable genes or prognostic biomarkers, enhancing our outlined observations. Disparities in endometrial cancer incidence and outcomes are multifactorial (39), necessitating pairing genetic associations with observations of histopathologic, molecular influences, and environmental exposures. This work builds a foundation for understanding genetic ancestry’s impact on somatic mutations and histologic subtypes in endometrial cancer, potentially explaining disparities.

## SUPPLEMENTARY FIGURES

**Supplementary Fig 1.**
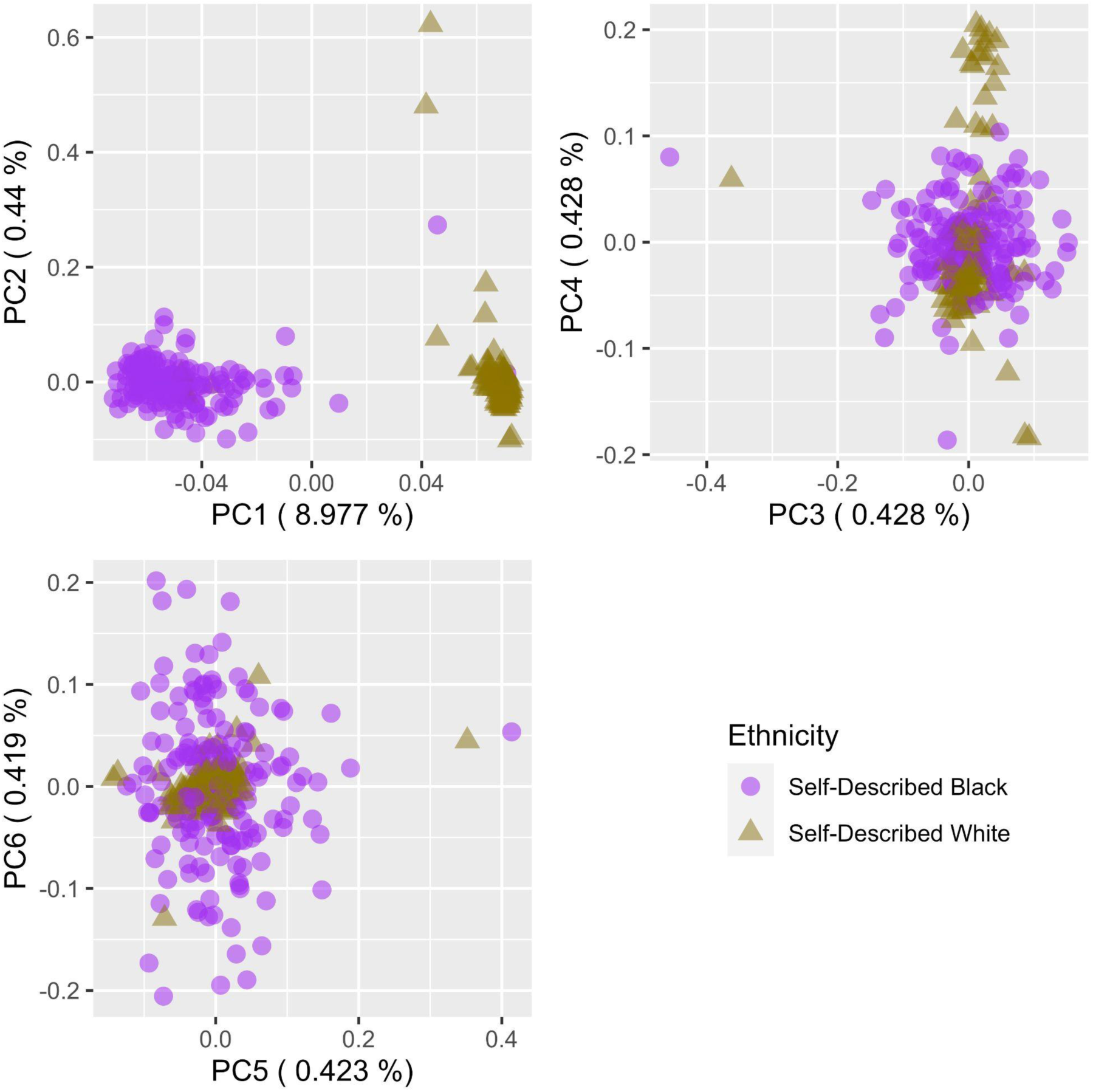
Principal componente analysis of germline whole-exome data. Dispersion plots show the relationsup between the first six principal componentes.

**Supplementary Fig 2.**
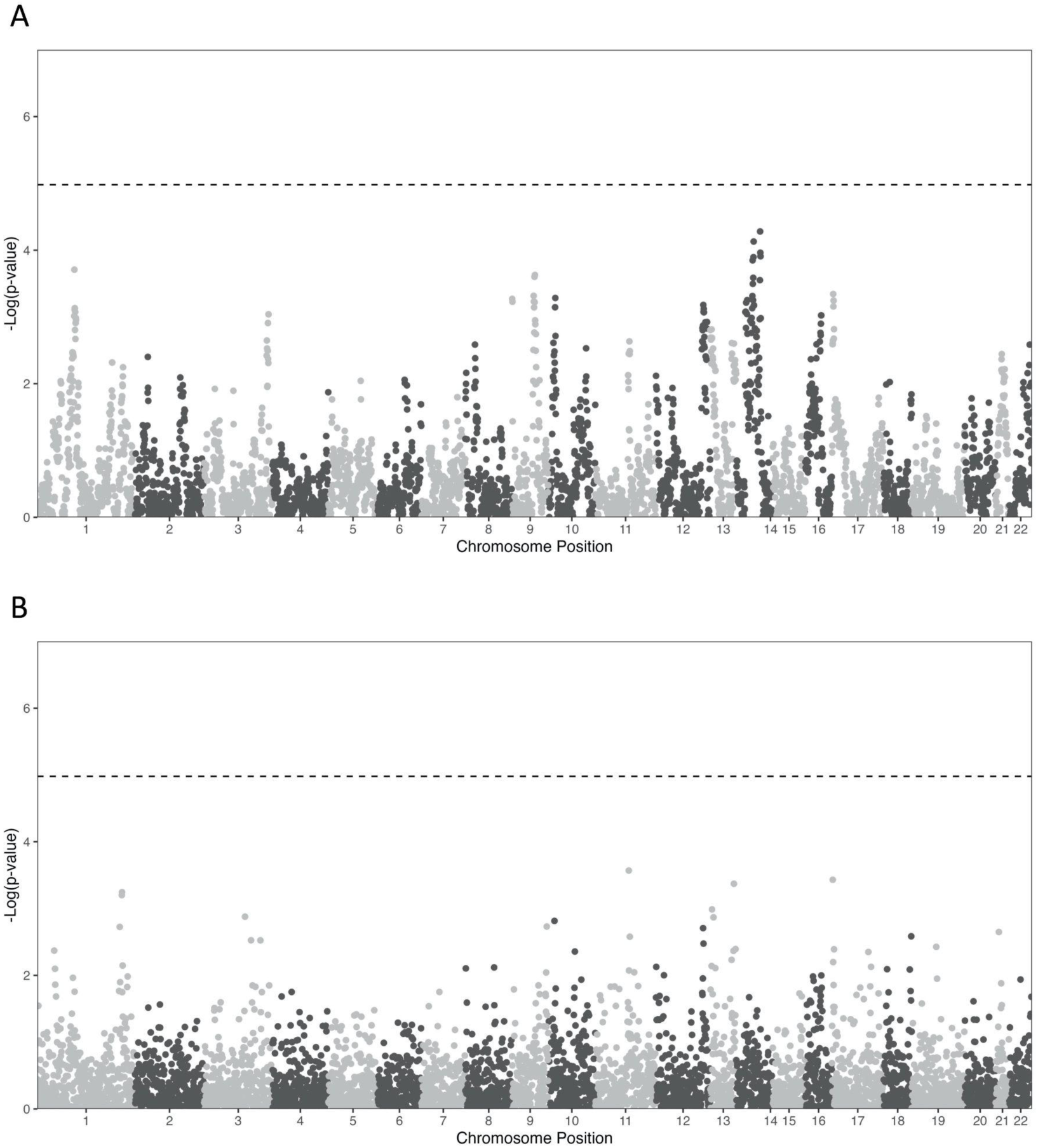
Admixture mapping model for the association between African local ancestry and somatic mutations. **(A)** Manhattan plot showing the outcome of the model examining the relationship between Africal local ancestry haplotypes per local ancestry regions and the total burden of somatic mutations genomewide. **(B)** Manhattan plot showing the outcome of the first admixture mapping-like model in which we look for correlations between the number of African local ancestry haplotypes per local ancestry region and the burden of somatic mutations (both synonymous and nonsynonymous) within that same region.

**Supplementary Figure 3.**
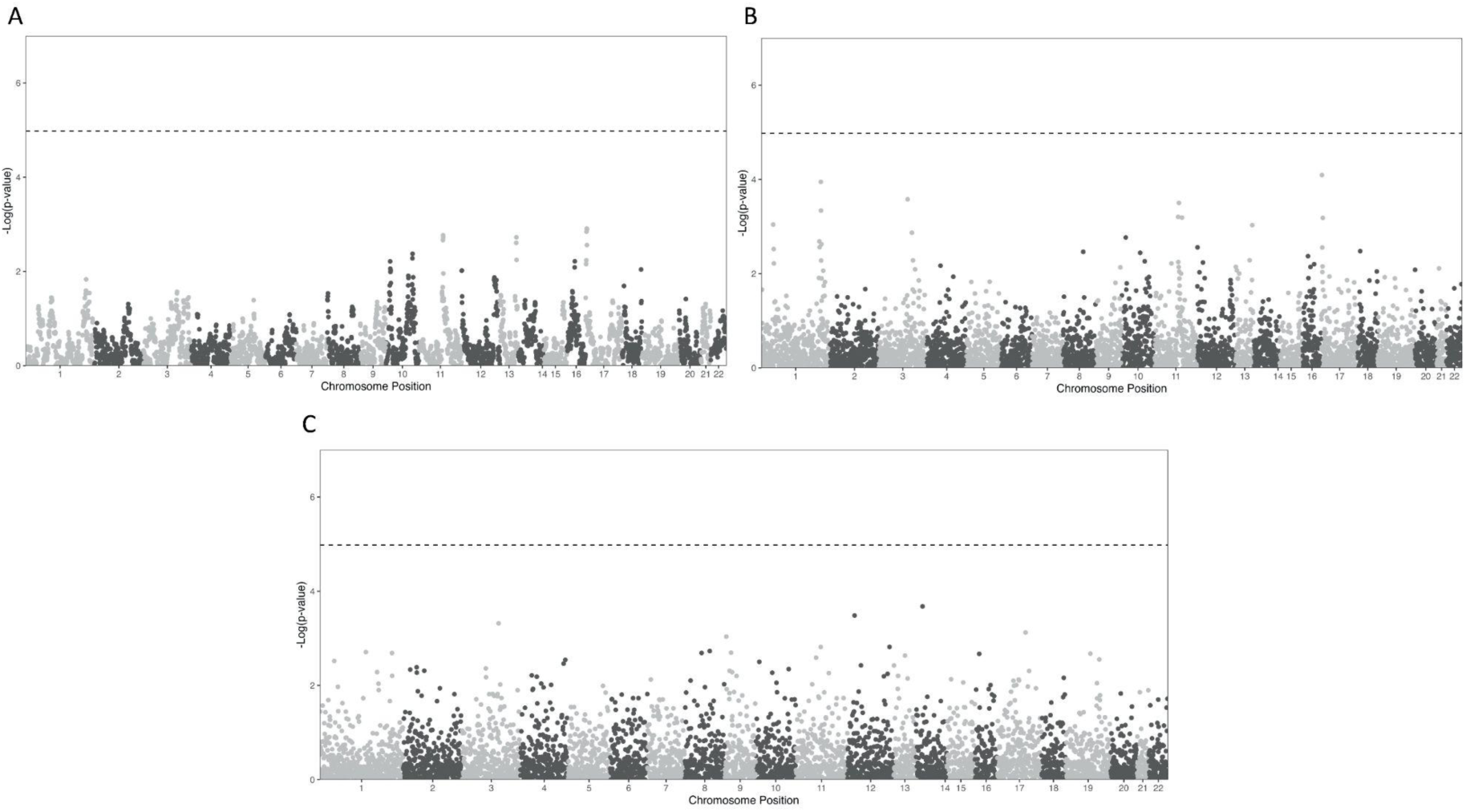
Admixture mapping-like models for the association between European local ancestry and somatic mutations. A) Association between European local ancestry haplotypes and the genome-wide frequency of all somatic mutations. B) Associations between European local ancestry haplotypes and local somatic mutation frequency. C) Same as in (A) but for nonsynonymous somatic mutation frequency, or TMB.

**Supplementary Fig 4.**
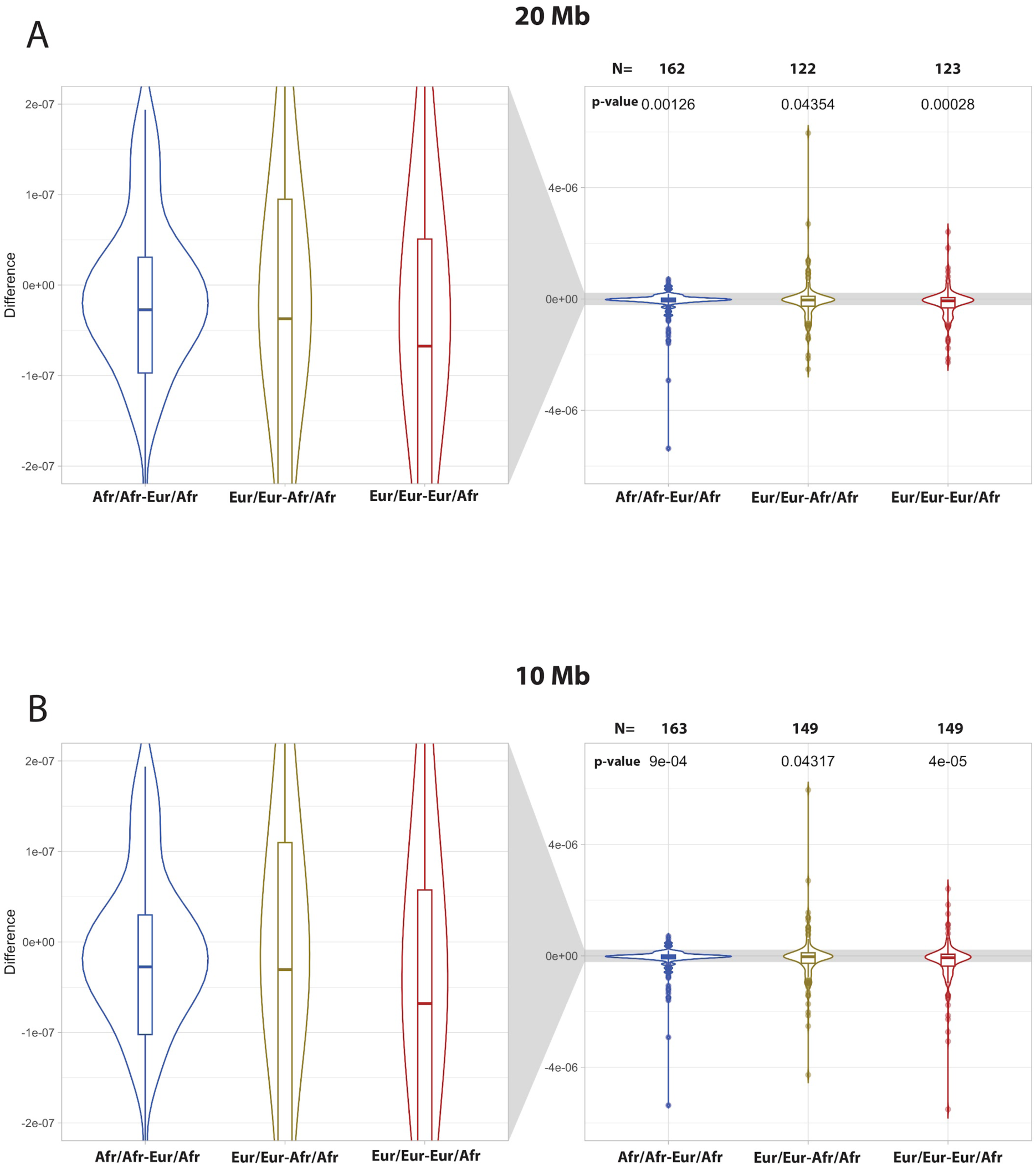
Relationship between Ancestral background of genomic regions and density of somatic mutations. Violin plots showing the distribution of the differences between pairs of piSD indexes. Each point represents the difference between diplotype rates in the same individual. We restricted the estimation of the differences to individuals harboring at least A) 20 and B) 10 Mb of each diplotype being compared. Significant differences are shown at the top of the plots.

**Supplementary Fig 5.**
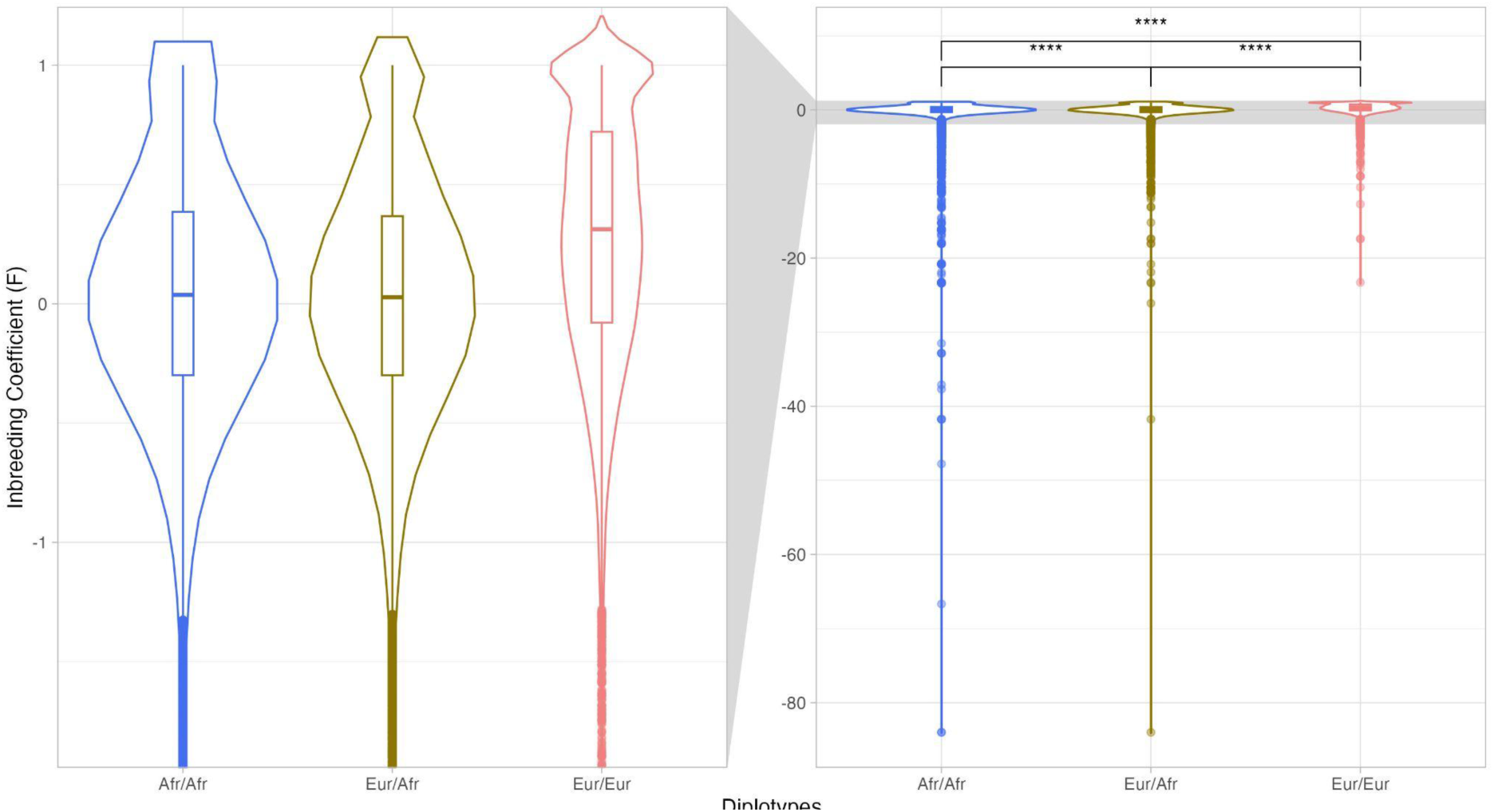
Relationship between inbreeding coefficient (F) and ancestral diplotypes. Each dot corresponds to a local ancestry window for an individual. On the left, boxplots show the relationship between F and the diplotypes. On the right zoomed area in the median region. Negative values of F indicate an excess of Heterozygosity.

**Supplementary Fig 6.**
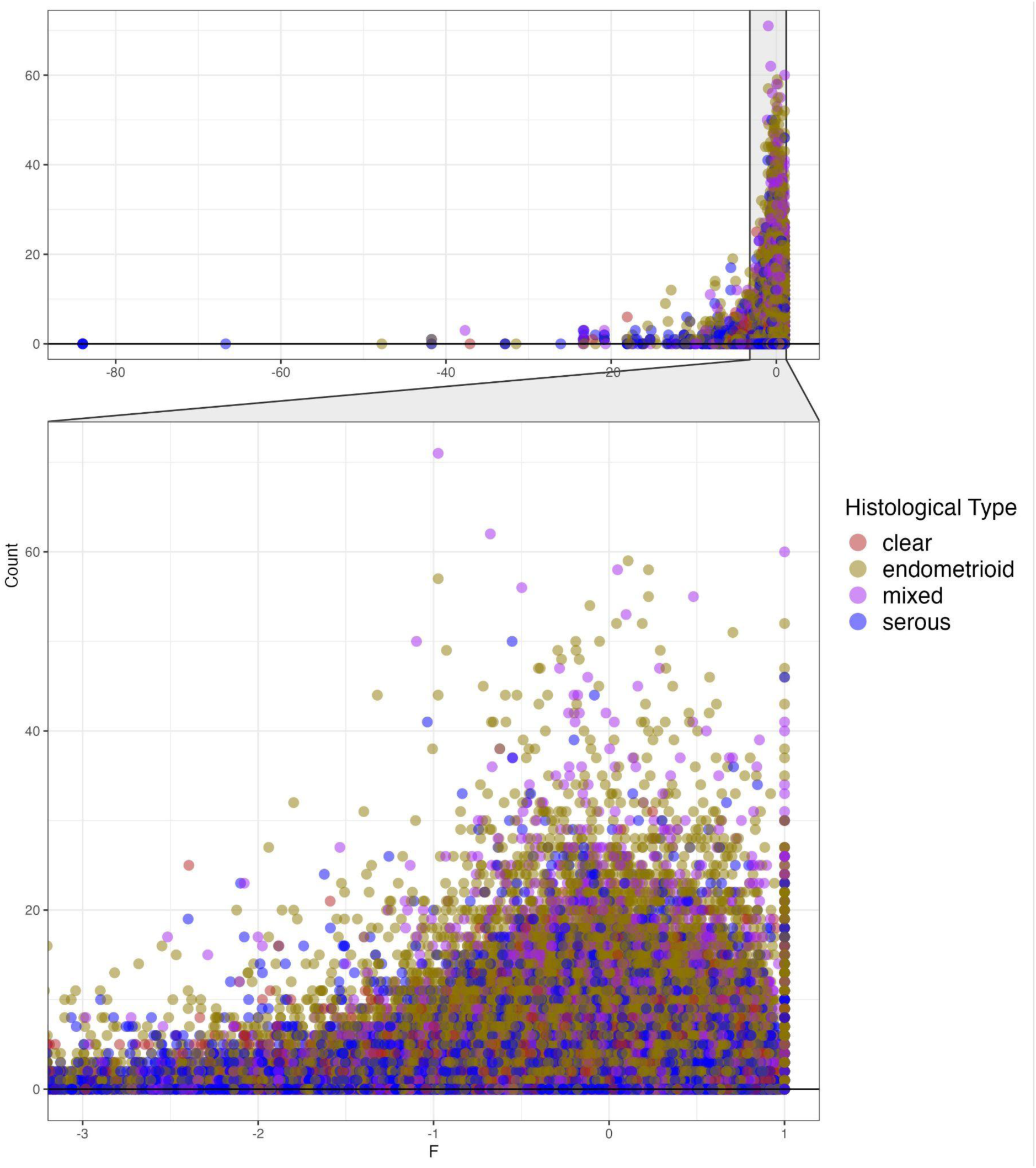
Relationship between Counts of Somatic mutations and inbreeding coefficient (F). Each dot corresponds to a size-specific window corresponding to local ancestry intervals in an individual. For each interval, we calculate the Somatic mutation counts and the F value. Negative F values indicate excess of heterozigosity. We also perform a negative binomial regression to explore the relationship between the Somatic counts and the F. A significant value and a negative beta was obtain for the inbreeding coefficient. Each point was colored based on the histological status (i.e., endometrioid, mixed, clear, and serous) of the tumor for each individual.

**Supplementary Fig 7.**
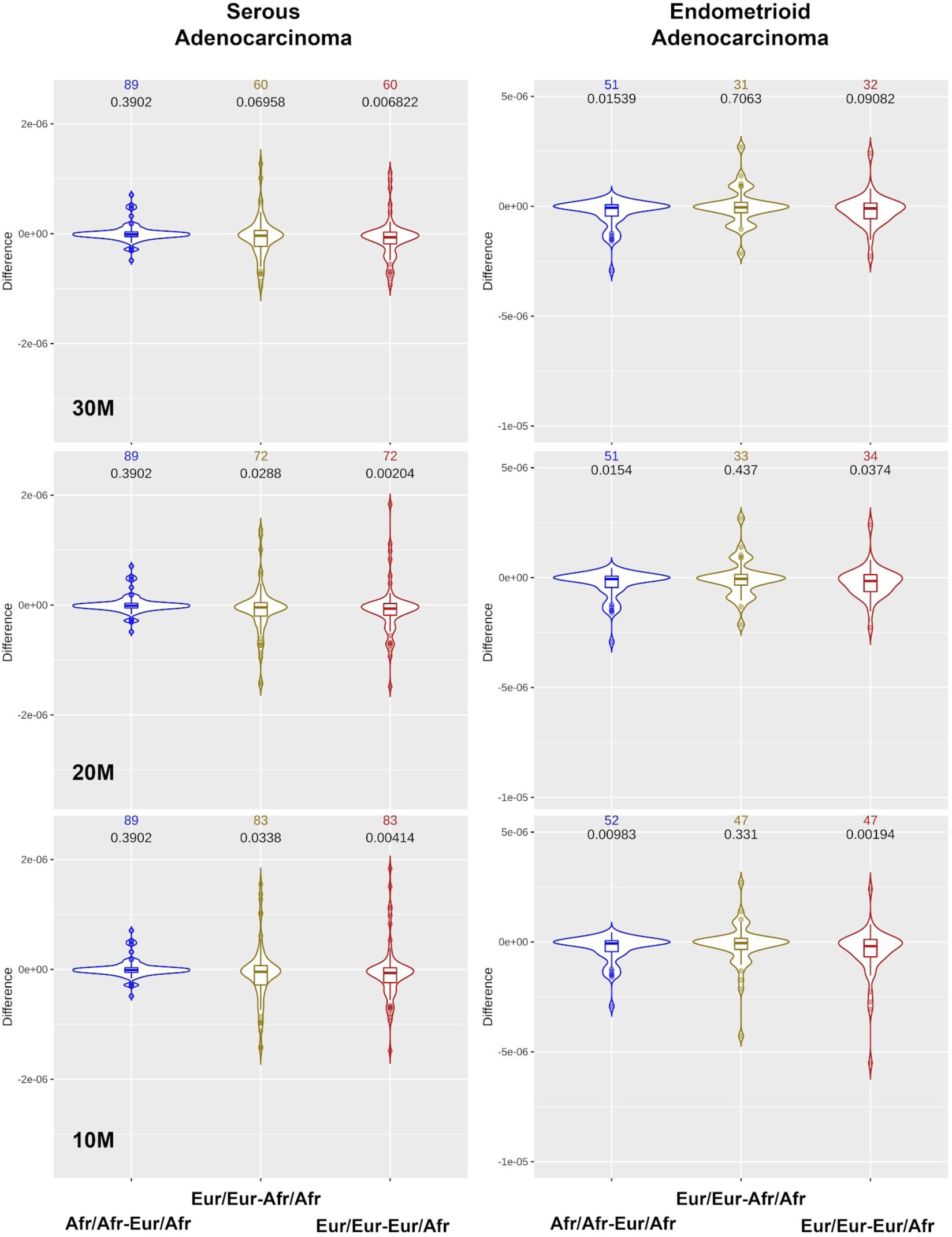
Relationship between Ancestral background of genomic regions and the density of somatic mutations. The ancestry of genomic regions was organized in terms of diplotypes. To calculate the difference between individual diplotype rates three thresholds were considered if the individual harbors at least 30, 20, and 10Mb of both diplotypes being compared. Violin plots showing the distribution of the differences between pairs of piSD indexes stratified per histological subtype. Significant differences are shown at the top of the plots.

**Supplementary Fig 8.**
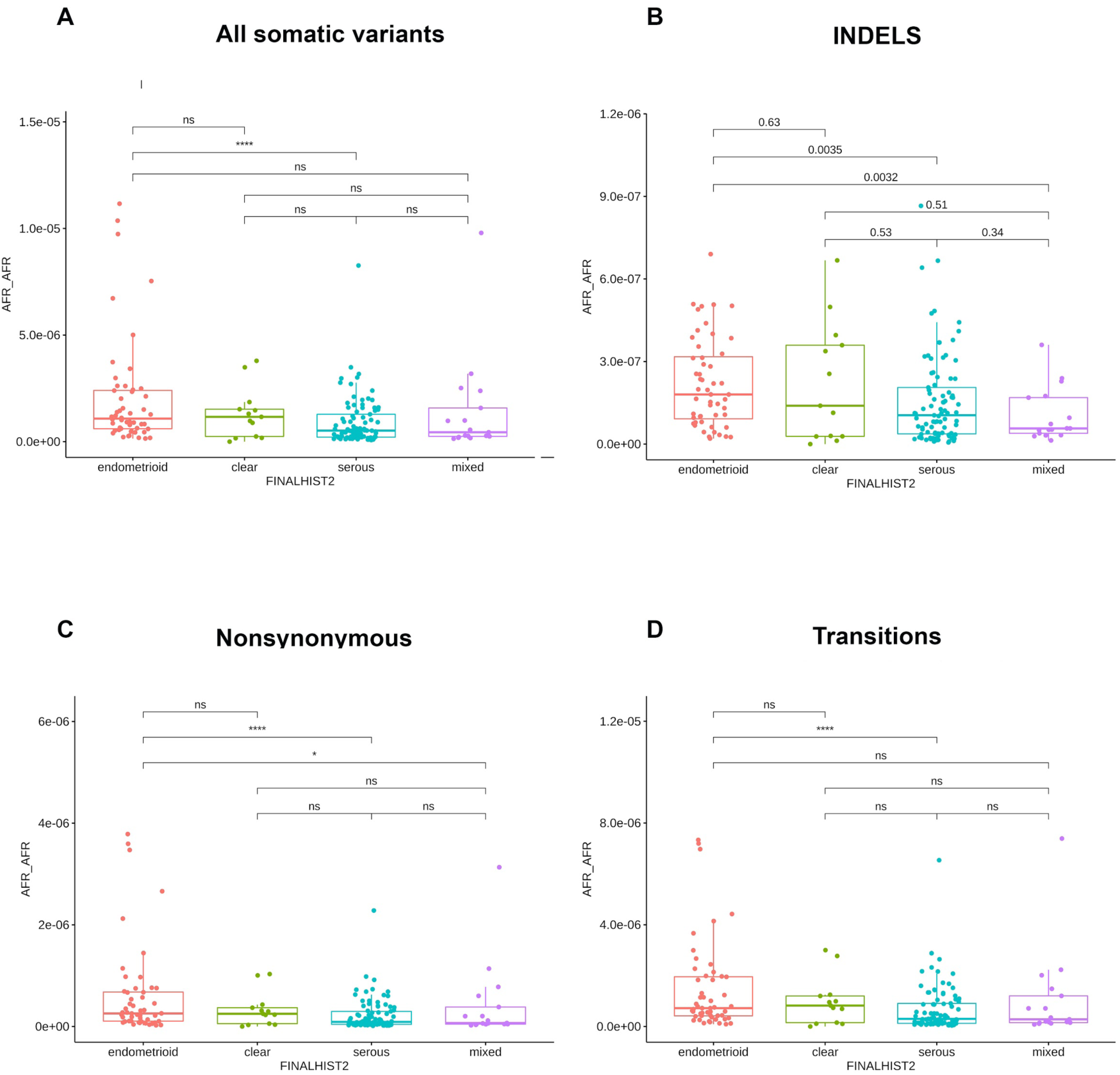
Average mutation burden per Mb observed in AFR/AFR diplotype per individual stratified per histological type and functional type of variants. The significant difference between distributions is indicated on the top of the box plots. ns: non-significant, *: (0.01, 0.05], **: (0.001, 0.01], ***: [0, 0.001].

**Supplementary Fig 9.**
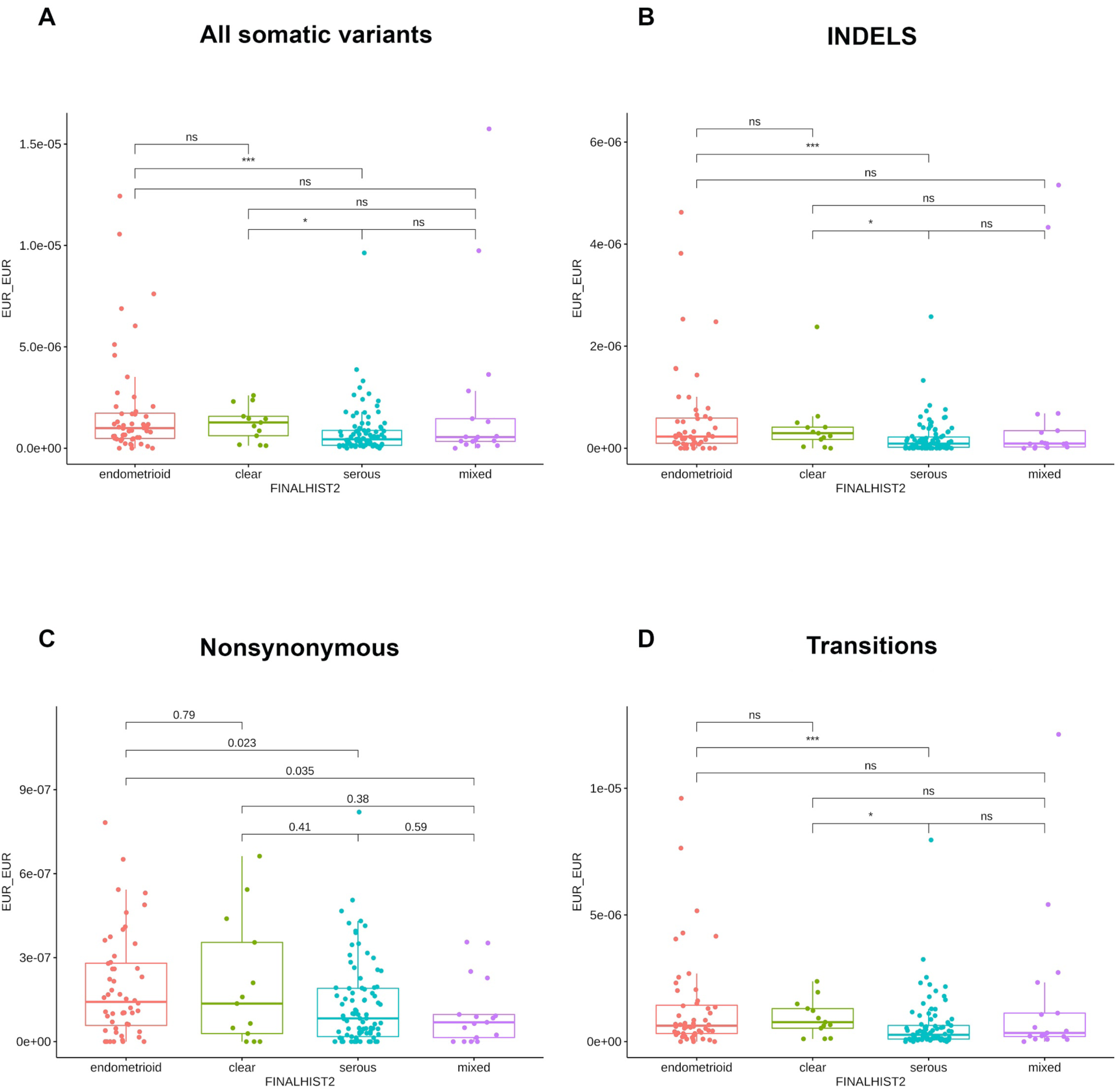
Average mutation burden per Mb observed in EUR/EUR diplotype per individual stratified per histological type and functional type of variants. The significant difference between distributions is indicated on the top of the box plots. ns: non-significant, *: (0.01, 0.05], **: (0.001, 0.01], ***: [0, 0.001].

**Supplementary Figure 10.**
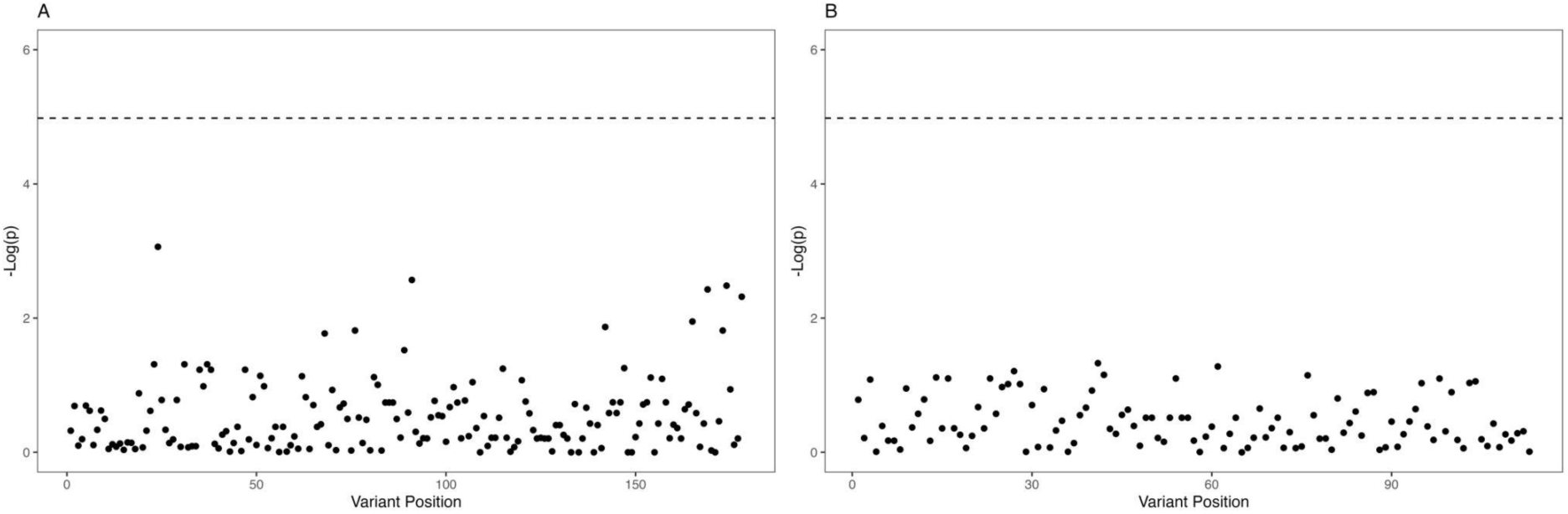
Firth Regression modeling association between variants and histological type within two identified regions. Regression within the regions A) chr9:112890610-113969108 B) chr9:131432751-131863024

## SUPPLEMENTARY TABLES

**Supplementary Table 1.** P-values for association between four global ancestry proportions (AFR:African, EUR:European, EAS:East Asian, NAT:Native American) and TMB for the self-described Black individuals.

| *ANCESTRY*<br>*LABEL* | *P-VALUE* |
| --- | --- |
| AFR | 0.379 |
| EUR | 0.433 |
| EAS | 0.556 |
| NAT | 0.582 |

**Supplementary Table 2.** Unstandardized and standardized coefficients for effect of significant regions noted in figure 1. Coefficients are expressed in mutation/Mb. Shown in parentheses are the 95% confidence intervals associated with each coefficient. Also shown is the gene content of each region, obtained from GENCODE v.41 (40). Genes for which the annotation overlaps multiple regions are shown for all regions that contain portions of the gene coordinates.

| *POSITION* | *UNSTANDARDIZED*<br>*COEFFICIENTS: B (95%*<br>*CI)* | *STANDARDIZED*<br>*COEFFICIENTS: $\beta$ (95%CI)* | *GENE CONTENT* |
| --- | --- | --- | --- |
| CHR 1:<br>62,834,807-64,009,249 | $2.5164 \times 10^{-2}$<br>($1.4870 \times 10^{-2} - 3.5458 \times 10^{-2}$) | $1.9542 \times 10^{-2}$<br>($9.146 \times 10^{-3} - 2.9938 \times 10^{-2}$) | ATG4C<br>FOXD3<br>ALG6<br>ITGB3BP<br>PGM1<br>ROR1 |
| CHR 1:<br>64,009,249-64,691,780 | $2.3386 \times 10^{-2}$<br>($1.3023 \times 10^{-2} - 3.3750 \times 10^{-2}$) | $1.7690 \times 10^{-2}$<br>($7.245 \times 10^{-3} - 2.8135 \times 10^{-2}$) | ROR1<br>UBE2U<br>CACHD1 |
| CHR 14:<br>81,533,807-87,934,009 | $2.2376 \times 10^{-2}$<br>($1.2598 \times 10^{-2} - 3.2154 \times 10^{-2}$) | $2.2376 \times 10^{-2}$<br>($1.2599 \times 10^{-2} - 3.2154 \times 10^{-2}$) | SEL1L<br>FLRT2<br>GALC |

